## Supplementary Information for "A hybrid stochastic-deterministic approach to explore multiple infection and evolution in HIV"

**Jesse Kreger[a], Natalia L. Komarova[a] and Dominik Wodarz[b,a]**

a) Department of Mathematics, University of California Irvine, Irvine CA 92697

b) Department of Population Health and Disease Prevention Program in Public Health
Susan and Henry Samueli College of Health Sciences, University of California, Irvine CA 92697

---

**3 Comparison of the ODE and stochastic/hybrid simulations in the context of infection dynamics** **6**

**4 Impact of multiple infection and synaptic transmission on mutant evolution: additional results** **10**

### 1 Modeling details

#### 1.1 Full description of mathematical model with different mutant strains and mutations

Mutation is an important component of HIV-1 infection and evolution. Therefore, we generalize the basic ODE model given in the main text by Equations (1)-(5) to include an arbitrary number of mutant strains and arbitrary fitness landscape.

We assume that forward and back point mutations can occur independently at $k$ locations, all with rate $\mu$. This creates $2^k$ viral strains. In the case $k = 3$ and by denoting the “unmutated” condition with a lowercase letter and the “mutated” condition with a capital letter, we have the wild-type (abc), three strains with a single point mutation (Abc, aBc, and abC), three strains with a double point mutation (ABc, AbB, and aBC), and a triple mutant strain (ABC). Upon each infection event, mutation can happen at each location. Therefore, any strain can mutate into any other strain, and the probability of such an event is based on the number of point mutations in which the two strains differ. The likelihood a strain mutates at $i \in \{0, 1, \ldots, k\}$ specified locations is $\mu^i(1-\mu)^{k-i}$. Define $D(\mathcal{S}_i, \mathcal{S}_j) \in \{0, 1, \ldots, k\}$ as the number of point mutations where strain $\mathcal{S}_i$ and $\mathcal{S}_j$ differ. The probability that strain $\mathcal{S}_i$ mutates into strain $\mathcal{S}_j$ (or vice versa) is then $\mu_{D(\mathcal{S}_i,\mathcal{S}_j)} := \mu^{D(\mathcal{S}_i,\mathcal{S}_j)}(1-\mu)^{k-D(\mathcal{S}_i,\mathcal{S}_j)}$. We denote the fitness of each strain by $F_i$ for $i = 1, \ldots, 2^k$.

Assume that we have two strains ($k = 1$), the wild-type and mutant. In order to model synaptic transmission with multiple strains and fitness considerations, we start by considering an infecting cell that contains $n$ wild-type viruses and $m$ mutant viruses, where $0 < n + m \leq N$. We denote the fitness of the wild-type as $F_1$ and fitness of mutant as $F_2$, where these parameters have the meaning of the probability of successful infection, i.e. $0 \leq F_1, F_2 \leq 1$. Here $F_2$ could be smaller (disadvantageous mutant), equal (neutral mutant), or larger (advantageous mutant) than $F_1$. Let us denote the fraction of wild-type and mutant viruses as

$$\nu = \frac{n}{n+m}, \quad \psi = \frac{m}{n+m},$$

respectively. Synaptic transmission is modeled as follows. We fix the number of viruses that are picked up for a synaptic transmission event, $S = 3$ (free virus transmission is similar, only with $S = 1$). Then, the following procedure is repeated $S$ times: a virus is selected from the infecting cell with the probability equal to its abundance in the cell (that is, wild-type viruses are picked with probability $\nu$ and mutants with probability $\psi$). Each virus that is picked will proceed to infect the target cell successfully with the probability given by its fitness (that is, $F_1$ for the wild-type and $F_2$ for the mutant). Each "pick" can result in three possibilities:

1. A wild-type virus will go on to be successful in infecting the target cell; this happens with probability $p_1 = \nu F_1$. We denote that by $*$ below.

2. A mutant virus will go on to be successful in infecting the target cell; this happens with probability $p_2 = \psi F_2$. We denote that by $X$ below.

3. An unsuccessful infection event, which happens with probability $p_3 = \nu(1 - F_1) + \psi(1 - F_2)$. We denote that by 0 below.

Therefore, under $S = 3$, a single synaptic transmission event can result in ten different infection events. Four of them ($***, **X, *XX, XXX$) result in an infection of the target cell with all $S = 3$ viruses (and these are the only events if $F_1 = F_2 = 1$). The other six events ($**0, *X0, XX0, *00, X00, 000$) result in an infection event with fewer than $S$ viruses. The probabilities of these events can be calculated by using multinomial distributions. In particular, given that the infecting cell is characterized by $(n, m)$, the probability of an event where $\hat{s}_1$ wild-type viruses and $\hat{s}_2$ mutant viruses go on to successfully infect the target cell is given by

$$P_{n,m}(\hat{s}_1, \hat{s}_2) = \frac{S!}{\hat{s}_1! \hat{s}_2! (S - \hat{s}_1 - \hat{s}_2)!} p_1^{\hat{s}_1} p_2^{\hat{s}_2} p_3^{S - \hat{s}_1 - \hat{s}_2}. \tag{1}$$

Note the following special cases. If the target cell has $n = 0$ (that is, it is only infected by the mutant), then $\nu = 1$ and $p_2 = F_2$. The only event with $S$ successful infections is $XXX$ and it happens with probability $F_2^S$. On the other hand, if $m = 0$, we have event $***$ with probability $F_1^S$. In other words, fitness properties of viruses are not erased if they are in cells that are not coinfected with both virus strains.

Next, we include the process of mutations. We assume that a virus can mutate upon entering the target cell, such that the process of mutation does not affect the success of infection. As there are only two strains, denote the probability that a wild-type virus mutates by $\mu$ and the probability that a mutant back-mutates to revert to a wild-type also by $\mu$. Let us suppose that a synaptic transmission event involves $\hat{s}_1$ wild-type and $\hat{s}_2$ mutant viruses, and consider the probability that upon entering the cell, we have $\hat{i}$ wild-type and $\hat{j}$ mutant viruses, where the change is due to mutations. We denote this probability as $Q_{\hat{s}_1;\hat{i},\hat{j}}$ (note that $\hat{s}_1 + \hat{s}_2 = \hat{i} + \hat{j}$). Suppose $\hat{a}$ out of $\hat{s}_1$ wild-type viruses mutate and $\hat{b}$ out of $\hat{s}_2$ viruses back-mutate. Then the number of (wild-type, mutant) viruses is $(\hat{s}_1 - \hat{a} + \hat{b}, \hat{s}_2 - \hat{b} + \hat{a}) = (\hat{\imath}, \hat{\jmath})$. Setting $\hat{a} = \hat{s}_1 - \hat{\imath} + \hat{b}$, we obtain

$$
\begin{aligned}
Q_{\hat{s}_1;\hat{\imath},\hat{\jmath}} &= \sum_{\hat{b}=0}^{\hat{s}_2} \frac{\hat{s}_1!}{\hat{a}!(\hat{s}_1-\hat{a})!}\mu^{\hat{a}}(1-\mu)^{\hat{s}_1-\hat{a}} \frac{\hat{s}_2!}{\hat{b}!(\hat{s}_2-\hat{b})!}\mu^{\hat{b}}(1-\mu)^{\hat{s}_2-\hat{b}} \\
&= \frac{\hat{s}_1!}{\hat{\imath}!}(1-\mu)^{\hat{\imath}}(1-\mu)^{\hat{\imath}+\hat{\jmath}-\hat{s}_1}\mu^{\hat{s}_1-\hat{\imath}}{}_2\tilde{F}_1\left(-\hat{\imath}, \hat{s}_1-\hat{\imath}-\hat{\jmath}, 1+\hat{s}_1-\hat{\imath}; \frac{\mu^2}{(1-\mu)^2}\right), \qquad (2)
\end{aligned}
$$

where we used the regularized hypergeometric function ${}_2\tilde{F}_1$.

For the general case when any number of viruses up to general $S$ can be transmitted successfully by synaptic transmission, we have that the full model with two virus strains is

$$
\begin{aligned}
\dot{x}_{0,0} &= \lambda - \beta x_{0,0}(Z_1+Z_2) - \gamma x_{0,0}\Big[\sum_{\hat{\imath}+\hat{\jmath}\le S}\ \sum_{0<n+m\le N}\ \sum_{\hat{s}_1=0}^{\hat{\imath}+\hat{\jmath}} P_{n,m}(\hat{s}_1, \hat{\imath}+\hat{\jmath}-\hat{s}_1)x_{n,m}\Big] - dx_{0,0}, \qquad (3) \\
\dot{x}_{i,j} &= \beta\Big[\big((1-\mu)Z_1+\mu Z_2\big)x_{i-1,j} + \big(\mu Z_1 + (1-\mu)Z_2\big)x_{i,j-1} - (Z_1+Z_2)x_{i,j}\Big] \\
&+ \gamma\Big[\sum_{\hat{\imath}+\hat{\jmath}\le S}\ \sum_{0<n+m\le N}\ \sum_{\hat{s}_1=0}^{\hat{\imath}+\hat{\jmath}} P_{n,m}(\hat{s}_1, \hat{\imath}+\hat{\jmath}-\hat{s}_1)Q_{\hat{s}_1;\hat{\imath},\hat{\jmath}}x_{n,m}(x_{i-\hat{\imath},j-\hat{\jmath}} - x_{i,j})\Big] - ax_{i,j}, \qquad (4)
\end{aligned}
$$

where $Z_1 = F_1\sum_{0<i+j\le N}\frac{i}{i+j}$ and $Z_2 = F_2\sum_{0<i+j\le N}\frac{j}{i+j}$, and with the appropriate adjustments that any population with a negative index is 0 and cells cannot be infected with more than $N$ total copies of virus. A system with more virus strains can easily be created as a generalization of this.

The number of equations per model where mutation can happen at $k$ independent locations is $2^{-k}(N+1)\binom{N+2^k}{2^k-1}$. To see this, we note that there are $2^k$ virus strains. The number of ways to distribute $j$ viral copies into the $2^k$ strains is $\binom{j+2^k-1}{2^k-1}$. Since we allow $j \in 0,\dots,N$, we have $\sum_{j=0}^{N}\binom{j+2^k-1}{2^k-1} = 2^{-k}(N+1)\binom{N+2^k}{2^k-1} = \binom{N+2^k}{2^k}$.

In the case of only free virus transmission ($\gamma = 0$), the fitness parameters can be interpreted as factors that modulate the rate of infection $\beta$, and so we can allow any $F_1, F_2 \ge 0$.

If we let $x_0$ denote the number of uninfected cells and $Z$ denote the sum of all infected cell subpopulations, we have that this model has a virus free steady state,

$$
\begin{aligned}
x_0 &= \frac{\lambda}{d}, \qquad (5) \\
Z &= 0, \qquad (6)
\end{aligned}
$$

and an infection steady state,

$$
\begin{aligned}
x_0 &= \frac{a}{\beta F + \gamma(1-(1-F)^S)}, \qquad (7) \\
Z &= \frac{\lambda}{a} - \frac{d}{\beta F + \gamma(1-(1-F)^S)}. \qquad (8)
\end{aligned}
$$

The stability of these steady states depends on the basic reproductive ratio, $R_0 = \frac{\lambda\left(\beta F + \gamma(1-(1-F)^S)\right)}{ad}$. Again we have that if $R_0 < 1$ the virus free steady state is stable and if $R_0 > 1$ then the infection steady state is stable.

##### 1.2 Determination of model parameters

Based on experimental evidence, we estimate the kinetic parameters in the model. Assume for simplicity that all virus strains are neutral with $F = 1$. It follows that the basic reproductive ratio is $R_0 = \frac{\lambda(\beta+\gamma)}{ad}$, the infection steady state for infected cells is $\frac{\lambda}{a} - \frac{d}{(\beta+\gamma)}$, and the virus free steady state for uninfected cells is $\frac{\lambda}{d}$.

We have an infection steady state value of $3.1 \times 10^7$ infected cells [2]. Assuming that the death rate of infected cells is approximately 2.2 days [10], we assume that $a = 0.45$ days$^{-1}$. Further, we set $R_0 = 8$ [11, 8]. Finally, we introduce parameter $\eta$ such that we have a total body population of $10^\eta$ lymphocytes at virus free steady state. Then we have the system of equations

$$3.1 \times 10^7 = \frac{\lambda}{a} - \frac{d}{(\beta+\gamma)}, \tag{9}$$

$$8 = \frac{\lambda(\beta+\gamma)}{ad}, \tag{10}$$

$$10^\eta = \frac{\lambda}{d}. \tag{11}$$

These equations uniquely imply that $\lambda \approx 1.59 \times 10^7$ days$^{-1}$, $d = \lambda \times 10^{-\eta}$ days$^{-1}$, and $\beta + \gamma = 3.6 \times 10^{-\eta}$ days$^{-1}$. We assume $S = 3$. Further, we choose $\eta = 9$ [2, 3], and therefore have $d \approx 0.0159$ days$^{-1}$. For initial conditions, we assume uninfected steady state and a single infected cell (infected with the wild-type) [9], that is $x_{0,0,\ldots,0} = \frac{\lambda}{d}$ and $x_{1,0,\ldots,0} = 1$.

At these parameters, triple and further mutant strains are negligible at peak infection, that is on average less than half a cell is infected with these strains when the number of infected cells is maximized (around seven days into infection). After peak infection, the system will settle to an equilibrium. These patterns can be seen by running the model with triple mutation first to peak infection, and then many days past peak infection.

#### 2 Details of the hybrid method

##### 2.1 Description of algorithm

At any given time, let $\mathbf{V}_l$ and $\mathbf{V}_s$ be vectors containing the large and small cell populations. We can then define the reduced system $d\mathbf{V}_l/dt = \mathbf{F}_l(\mathbf{V}_l)$ derived from the full system by: (1) retaining only the equations for the large cell populations $\mathbf{V}_l$; (2) keeping constant the contributions of the small populations $\mathbf{V}_s$. If the components of $\mathbf{V}_l$ are sufficiently large, there will be a time interval $(t, t+\tau)$, where the deterministic solution of the reduced ODE will approximate the trajectories of the large populations in a stochastic implementation of the full system.

When an infection event occurs (either free virus transmission or cell-to-cell transmission), the target cell moves from its current subpopulation to its new subpopulation of higher multiplicity. We refer to the original cell population as the *minus cell subpopulation* (because it has lost a member), and to the new subpopulation as the *plus cell subpopulation* (because it has gained a member).

The events in the model are uninfected cell generation, cell death, and infection with up to $S$ copies of combinations of the $2^k$ viral strains. In Gillespie's method, every event $\nu$ has a given propensity $a_\nu(\mathbf{V})$. The time at which the next event $\nu$ will occur is exponentially distributed with intensity $a_\nu(\mathbf{V})$. In the hybrid approach, uninfected cell generation, death of large populations, and infection where both the minus cell population and plus cell population are large are modeled deterministically (using the reduced system), while cell death and infection of small populations are modeled stochastically, with propensities $a_\nu(\mathbf{V}_s, \mathbf{V}_l(t))$ that now vary continuously with time.

If the minus cell population and the plus cell population are of different size classification (one is small and the other is large), then infection between these two populations is modeled with a “half and half reaction” (see next Section). For further details on the hybrid algorithm, see [12].

#### 2.2 Classification of reactions

This is an explanation of how individual reactions are designated small or large. This is a system with $N = 4$, no mutation, and only free virus transmission. Let $Z(t) = \sum_{i=1}^{4} x_i(t)$ be the sum of all infected subpopulations. The equations in the absence of synaptic transmission are

$$\dot{x}_0 = \lambda - \beta Z x_0 - d x_0 \tag{12}$$

$$\dot{x}_1 = \beta Z x_0 \boldsymbol{- \beta Z x_1} - a x_1 \tag{13}$$

$$\dot{x}_2 = \boldsymbol{\beta Z x_1} - \beta Z x_2 - a x_2 \tag{14}$$

$$\dot{x}_3 = \beta Z (x_2 - x_3) - a x_3 \tag{15}$$

$$\dot{x}_4 = \beta Z (x_3) - a x_4 \tag{16}$$

Assume we are at time $t^*$ and we have size threshold $\mathcal{M}^*$. Assume that that $x_0(t^*) \geq \mathcal{M}^*$, $x_1(t^*) \geq \mathcal{M}^*$, $x_2(t^*) < \mathcal{M}^*$, $x_3(t^*) < \mathcal{M}^*$, and $x_4(t^*) < \mathcal{M}^*$. Then at time $t^*$ we have that populations $x_0$ and $x_1$ are large and all other populations are small.

Since $x_0$ and $x_1$ are large and all other populations are small, we only update (12) and (13) deterministically. Everything in Equations (14)-(16) is stochastic. Therefore, it makes sense that (i) uninfected cell generation is small/large if $x_0$ is small/large; (ii) death of population $x_i$ is small/large if $x_i$ is small/large; (iii) infection of population $x_i$ is small if $x_i$ (the minus population) or the plus population is small, and large otherwise.

Here, the reaction that infects population $x_1$ is updated deterministically in Equation (13) and also stochastically because population $x_2$ is small. Therefore, we lose a singly infected cell both deterministically and stochastically, and only gain an $x_2$ stochastically. Therefore, we designate the reaction that infects populations $x_1$ a “half and half” reaction. This means that $x_1$ is updated deterministically in Equation (13) and also stochastically because population $x_2$ is small, but during the stochastic reaction we only gain an $x_2$ and do not lose an $x_1$. So in essence we lose $x_1$ to infection deterministically, and gain $x_2$ by infection stochastically (but do not lose an $x_1$). More generally, we have a half and half reaction whenever the minus population is large and the plus population is small, or the minus population is small and the plus population is large.

In this system, since there is only free virus transmission the plus population is always $x_{i+1}$. However, in a system with synaptic cell-to-cell transmission and a mutant strain, the plus population can be $x_{i+1,j}$, $x_{i,j+1}$, $x_{i+S,j}$, $x_{i,j+S}$, etc. So under these assumptions, in the ODE system given by Equations (12)-(16), we have that all unbolded terms in Equations (12)-(13) are the large deterministic reactions, the **bolded** terms are the half and half reactions, and all other terms are the stochastic reactions.

#### 2.3 Choosing a size threshold $\mathcal{M}$

The rate of stochastic extinction of the hybrid method should match the rate of stochastic extinction of the simulations in the fully stochastic case [14, 1]. As described in the main text, we define $\hat{\mathcal{M}}$ as the lower bound on the size threshold $\mathcal{M}$. Therefore we have that

$$\hat{\mathcal{M}} = \lceil \ln\left(1 + \frac{R_0 - 1}{\delta R_0}\right) / \ln R_0 \rceil, \tag{17}$$

where $\lceil . \rceil$ denotes the ceiling function. Since increasing $R_0$ monotonically leads to a higher rate of successful infections, we see that increasing $R_0$ leads to decreasing $\hat{\mathcal{M}}$. Table 1 shows lower bound $\hat{\mathcal{M}}$ for a variety of $R_0 > 1$ and difference thresholds $\delta$.

When implementing the hybrid algorithm, the ODE step size $h$ must also be taken into account. The step size $h$ must be small enough such that i) in the completely deterministic case it gives an accurate numerical solution of the ODE solution, and ii) in the hybrid case the stochastic events and dynamics are captured. This only becomes an issue when size threshold $\mathcal{M}$ is very large (so there are many populations and $\tau$ becomes quite small for multiple reactions) but there are both small and large populations (so that some populations are still updating deterministically). In this case, the ODEs for the uninfected population becomes stiff and smaller step size $h$ might be needed to capture the initial decline of the uninfected cell population.

| $R_0$ | $\hat{\mathcal{M}}$ for $\delta = 10^{-4}$ | $\hat{\mathcal{M}}$ for $\delta = 10^{-5}$ | $\hat{\mathcal{M}}$ for $\delta = 10^{-6}$ |
|---|---|---|---|
| 1.001 | 2399 | 4617 | 6912 |
| 1.01 | 463 | 694 | 925 |
| 1.1 | 72 | 96 | 120 |
| 1.5 | 21 | 26 | 32 |
| 2 | 13 | 16 | 19 |
| 4 | 7 | 9 | 10 |
| 6 | 6 | 7 | 8 |
| 8 | 5 | 6 | 7 |

Table 1: $\hat{\mathcal{M}}$ values for a variety of $R_0 > 1$ and difference thresholds $\delta$. We define $\hat{\mathcal{M}}$ according to Equation (17).

### 3 Comparison of the ODE and stochastic/hybrid simulations in the context of infection dynamics

The deterministic ODEs do not provide information on the distribution of mutant numbers. The hybrid method is especially useful here, because it allows for efficient stochastic computations at large population sizes, and can generate mutant distributions rather than just averages.

Figure S1 shows an example of the distribution of the number of cells infected with a neutral single and double mutant strain of the virus when the infected cell population reaches $10^6$ cells for hybrid simulations with size threshold $\mathcal{M} = 50$ and $\mathcal{M} = 500$ and only free virus transmission. Here, we see that for $R_0 = 8$, size threshold $\mathcal{M} = 50$ is large enough to provide a good approximation of the fully stochastic dynamics. Figure S2 shows that for $R_0 = 1.5$, $\mathcal{M} = 50$ is not large enough to capture the stochastic dynamics, and larger size thresholds are needed. As discussed in the main text, lower $R_0$ is associated with larger stochastic effects.

Furthermore, an important benchmark of infection is the emergence of mutant strains of the virus. The hybrid method allows us to compute full distributions for when the first mutant is expected to appear stochastically (by a mutation from a wild-type virus). Figure S3 demonstrates how the deterministic model predictions for the first mutant virus to appear can differ from averages over many stochastic runs simulated using the computationally efficient hybrid algorithm.

Finally, while we have shown that ODEs cannot accurately describe the average behavior of the stochastic model, the hybrid method (with a sufficient size threshold) is able to do so. Figure

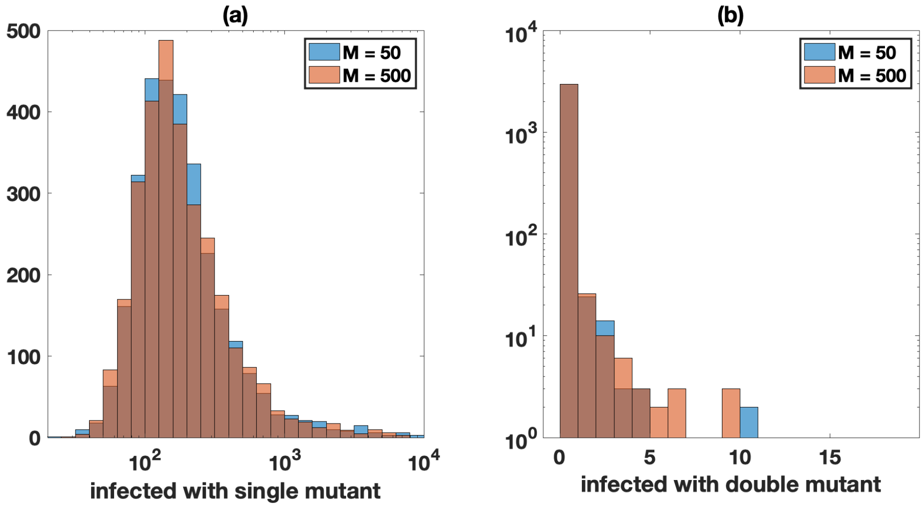

Figure S1: Distribution of the number of cells infected with the neutral single and double mutant strain of the virus (double mutant model, $F = 1$ for all strains) when the infected cell population reaches $10^6$ cells for hybrid simulations with size threshold $\mathcal{M} = 50$ and $\mathcal{M} = 500$. Simulations include only free virus transmission. For each size threshold $3 \times 10^3$ simulations were performed. (a) Number of cells infected with one of the single mutant strains. (b) Number of cells infected with the double mutant strain. The parameters are $N = 3$, $\mu = 3 \times 10^{-5}$, $\lambda = 1.59 \times 10^7$, $\beta = 3.60 \times 10^{-9}$, $\gamma = 0$, $a = 0.45$, $d = 0.016$, and $R_0 = 8$. For both single and double mutants, the probability distributions of the cell numbers infected with mutant viruses are not statistically different for $\mathcal{M} = 50$ and $\mathcal{M} = 500$ ($p > 0.1$ by Kolmogorov-Smirnov test).

S4(a) compares the number of cells infected with the mutant at infected population size $10^4$ and Figure S4(b) compares the time at first mutant generation for simulations in which all infected cell populations are always treated stochastically versus simulations with $\mathcal{M} = 50$. In both cases, the Kolmogorov-Smirnov test between the two distributions does not suggest that there is a statistically significant difference.

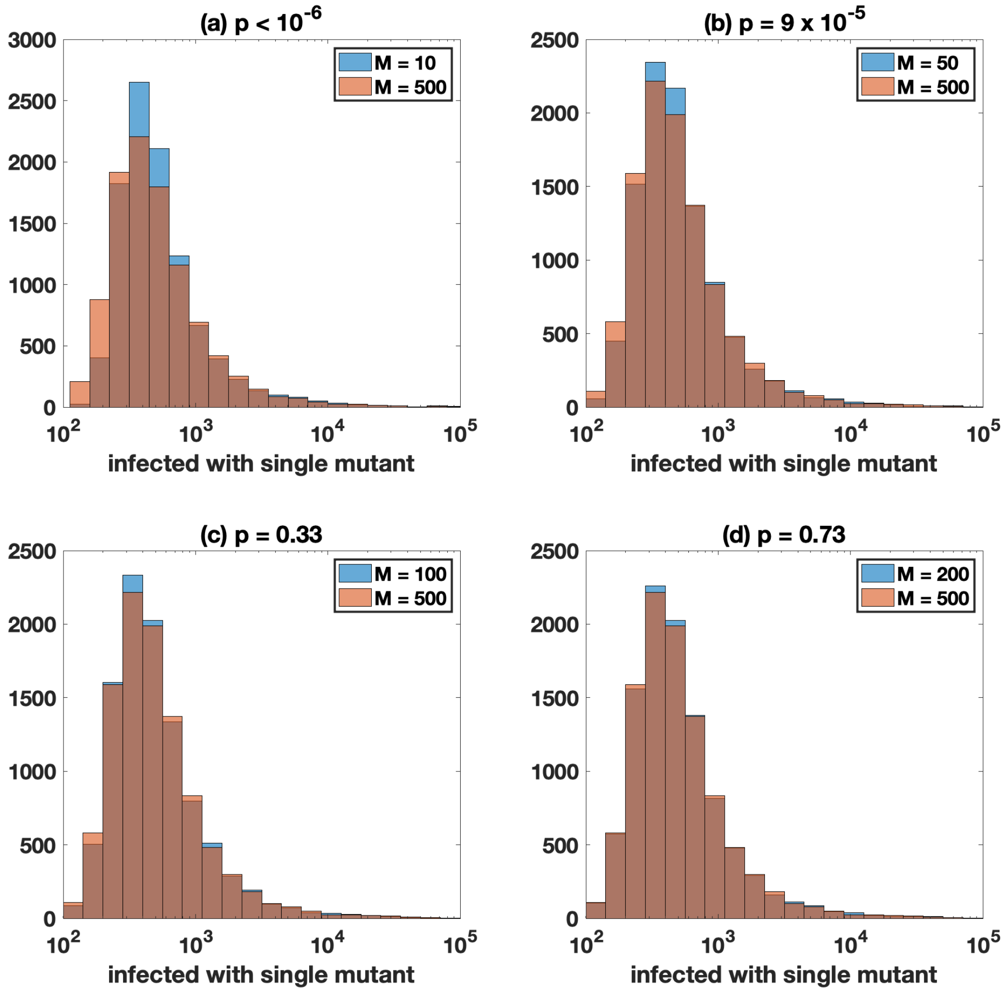

Figure S2: Distribution of the number of cells infected with the neutral single mutant strain of the virus (single mutant model, $F_{\text{wild-type}} = F_{\text{mutant}} = 1$) when the infected cell population reaches $10^6$ cells for hybrid simulations with only free virus transmission. Distributions for each size threshold represent $10^4$ simulations. The $p$-value from Kolmogorov-Smirnov test is shown for each comparison. (a) Size threshold $\mathcal{M} = 10$ and $\mathcal{M} = 500$. (b) Size threshold $\mathcal{M} = 50$ and $\mathcal{M} = 500$. (c) Size threshold $\mathcal{M} = 100$ and $\mathcal{M} = 500$. (d) Size threshold $\mathcal{M} = 200$ and $\mathcal{M} = 500$. Here $N = 1$ and the other parameters are $\mu = 3 \times 10^{-5}$, $\lambda = 1.59 \times 10^7$, $\beta = 3.60 \times 10^{-9}$, $\gamma = 0$, $d = 0.016$, and $R_0 = 1.5$.

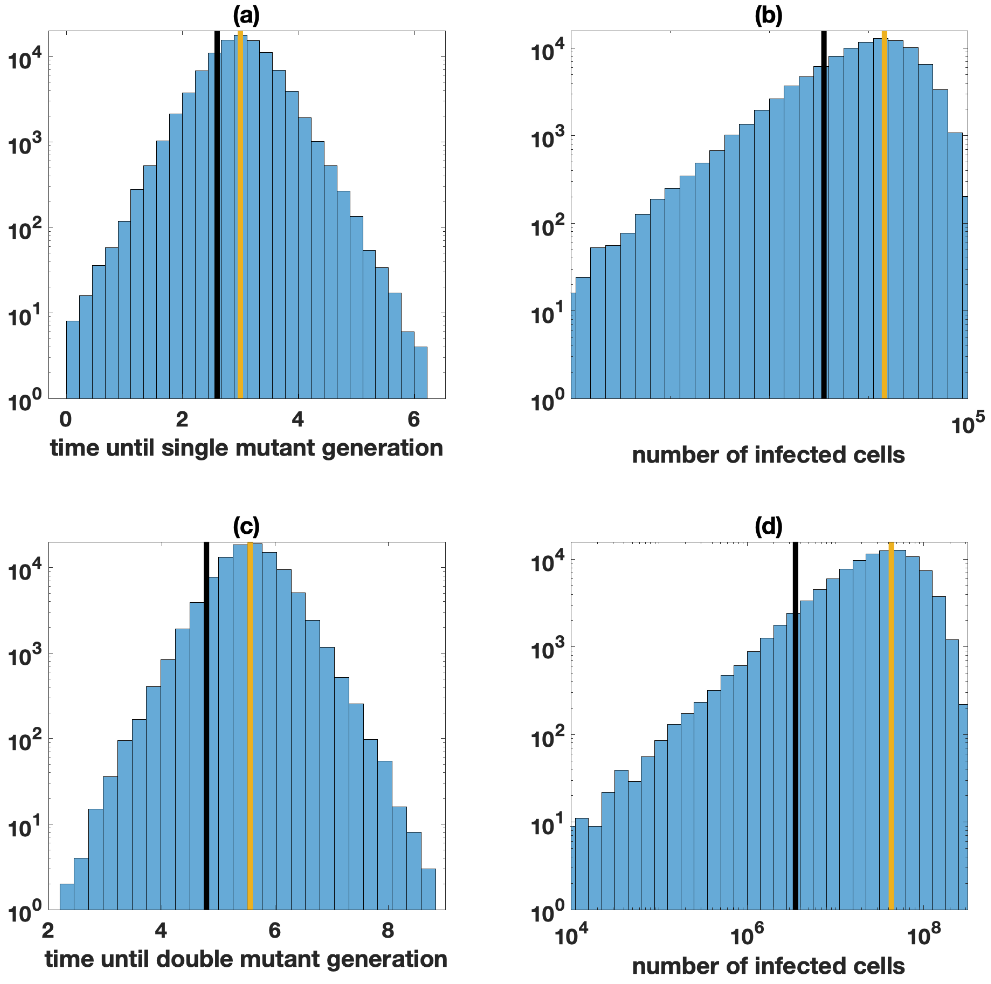

Figure S3: Distribution of generation time of first single/double mutant virus in the double mutation model in the context of only free virus transmission (with all strains neutral). Simulations in which infections are not established are discarded when calculating the averages. The deterministic prediction is denoted with the black vertical line and the hybrid average is denoted with the yellow vertical line. Distributions represent $10^5$ hybrid simulations with size threshold $\mathcal{M} = 50$. (a) Time until either single mutant generation. The deterministic prediction that the first single mutant virus (of both strains) will be generated is around 2.6 days, whereas in the stochastic case it is around 3 days. (b) Number of infected cells at first single mutant generation. The deterministic prediction is that the number of infected cells is around $3.5 \times 10^3$, whereas in the stochastic case it is around $1.4 \times 10^4$. (c) Time until double mutant generation. The deterministic prediction that the first double mutant virus will be generated is around 4.8 days, whereas in the stochastic case it is around 5.6 days. (d) Number of infected cells at first double mutant generation. The deterministic prediction is that the number of infected cells is around $3.5 \times 10^6$, whereas in the stochastic case it is around $4.3 \times 10^7$. The parameters are $N = 3,\ \mu = 3 \times 10^{-5},\ \lambda = 1.59 \times 10^7,\ \beta = 3.60 \times 10^{-9},\ \gamma = 0,\ a = 0.45,\ d = 0.016$, and $R_0 = 8$.

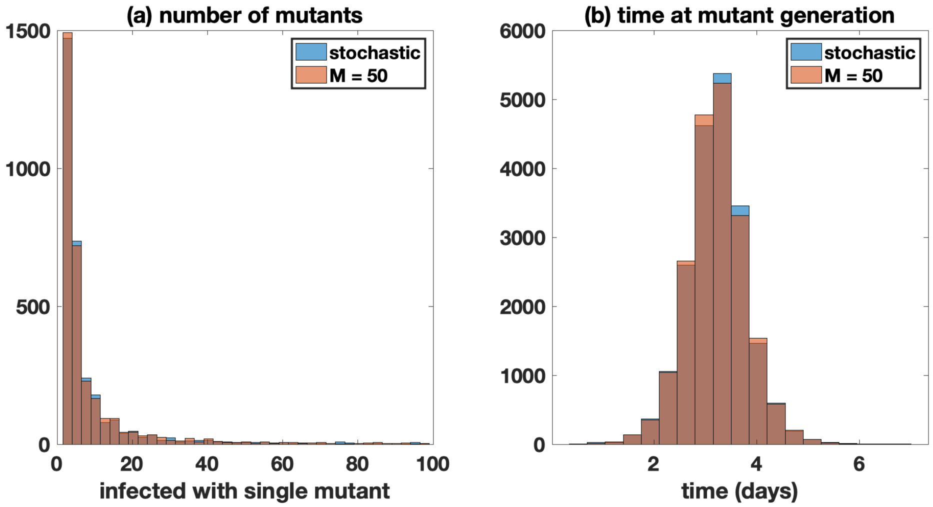

Figure S4: Comparing hybrid simulations with $\mathcal{M} = 50$ (red) with simulations in which the infected subpopulations are always treated stochastically (blue). Simulations in which infections are not established are discarded when calculating the averages. Distributions represent $2 \times 10^4$ simulations. (a) The number of cells infected with the mutant at infected population size $10^4$. (b) The time at first mutant generation. The parameters are $F_{\text{wild-type}} = F_{\text{mutant}} = 1,\ N = 3,\ \mu = 3 \times 10^{-5},\ \lambda = 1.59 \times 10^7,\ \beta = 3.60 \times 10^{-9},\ \gamma = 0,\ a = 0.45,\ d = 0.016$, and $R_0 = 8$.

### 4 Impact of multiple infection and synaptic transmission on mutant evolution: additional results

We consider the effect of multiple infection on mutant dynamics, as well as different scenarios where fitness is dependent upon the contents of a cell (i.e. complementation, interference). Thus, the multiplicity of infection is important. Widely varying estimates for average infection multiplicities have been published [6, 4, 5, 13], and there is some uncertainty. Figure S5 shows the deterministic time series for an infection with a single neutral mutant strain in the presence of only free virus or only synaptic transmission. For only free virus transmission, the multiplicity of infection near peak infection is 2 and reaches a maximum of 6 just after peak infection. For only synaptic transmission, the multiplicity of infection near peak infection is 7 and reaches a maximum of 17 just after peak infection. Stochastic simulations give similar values for the multiplicity of infection, and are very consistent with standard deviations usually less than $10^{-2}$. Therefore, as we increase the contribution of synaptic transmission, the number of coinfected cells increases, which leads to more pronounced effects of interactions between different virus strains within cells such as complementation and interference.

We can also use the hybrid method to determine the distribution of multiplicity of infection near peak infection, based on different transmission strategies. Figure S6 shows the distribution of the number of cells infected with different numbers of viral copies on average, for only free virus transmission (Figure S6(a)) and half free virus transmission and half synaptic transmission (Figure S6(b)). The distribution for only synaptic transmission will be the same as only free virus transmission, except the number of viral copies (horizontal axis) is scaled by $S$ (i.e. $1 \to S,\ 2 \to 2S$) and all other values will contain no cells. As done in [7], the steady states for each population can be determined analytically.

Mutant evolution is impacted by multiple infection [15]. Figure S7 shows the distributions of cells infected with neutral single and double mutants in the presence and absence of multiple infection at relatively low virus loads. Here, the average number of mutants is the same, whether multiple infection is assumed to occur or not because of the low multiplicity of infection at lower

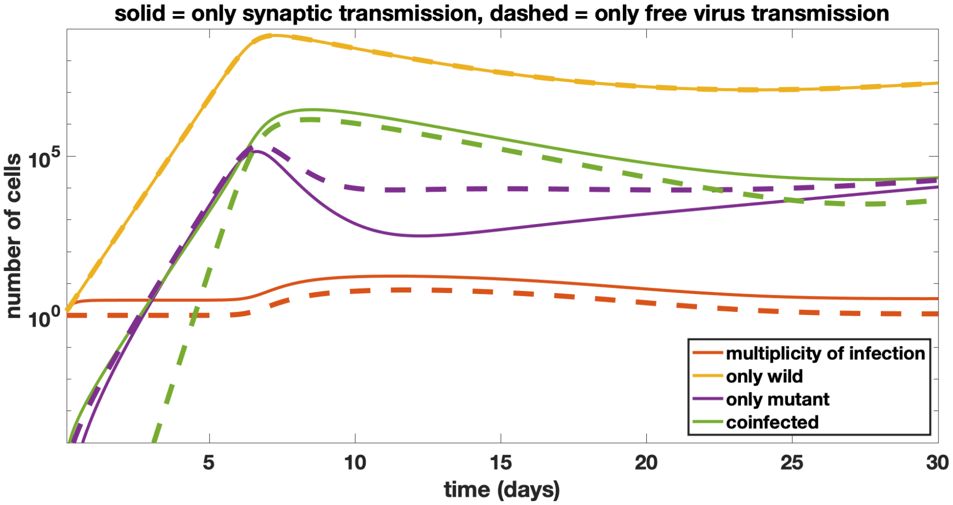

Figure S5: Deterministic time series evolution of an infection with a neutral mutant strain with only free virus (dashed lines, $\gamma = 0$) or only synaptic transmission (solid lines, $\beta = 0$). Parameters are $N = 25$, $\beta + \gamma = c = 3.6 \times 10^{-9}$, $\mu = 3 \times 10^{-5}$, $\lambda = 1.59 \times 10^7$, $a = 0.45$, and $d = 0.016$. The multiplicity of infection is shown with the red lines, the number of cells infected with only the wild-type virus are shown with the yellow lines, the number of cells infected with only the mutant are shown with the purple lines, and the number of cells coinfected with both the wild-type and mutant are shown with the green lines.

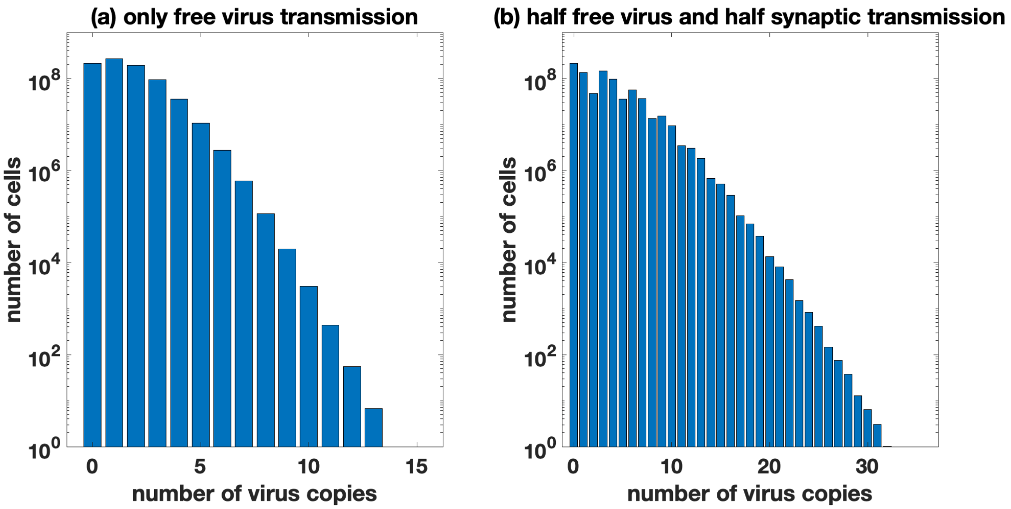

Figure S6: Distribution of the multiplicity of infection near peak infection. (a) Histograms for the average number of cells infected with the given number of viral copies for only free virus transmission. (b) Histograms for the average number of cells infected with the given number of viral copies for half free virus transmission and half synaptic transmission. The horizontal axis is the number of virus copies and the vertical axis is the average number of cells that are infected with that number of viral copies near peak infection. Infected with zero copies corresponds to the uninfected cells. Distributions were averaged over $5 \times 10^2$ hybrid simulations with size threshold $\mathcal{M} = 50$. Simulations are stopped when the infected cell population is close to peak infection ($6 \times 10^8$ cells). Parameters are $\beta + \gamma = c = 3.6 \times 10^{-9}$, $\mu = 3 \times 10^{-5}$, $\lambda = 1.59 \times 10^7$, $a = 0.45$, and $d = 0.016$, except that maximum multiplicity of infection $N$ is set to be large enough such that no cells reach this threshold.

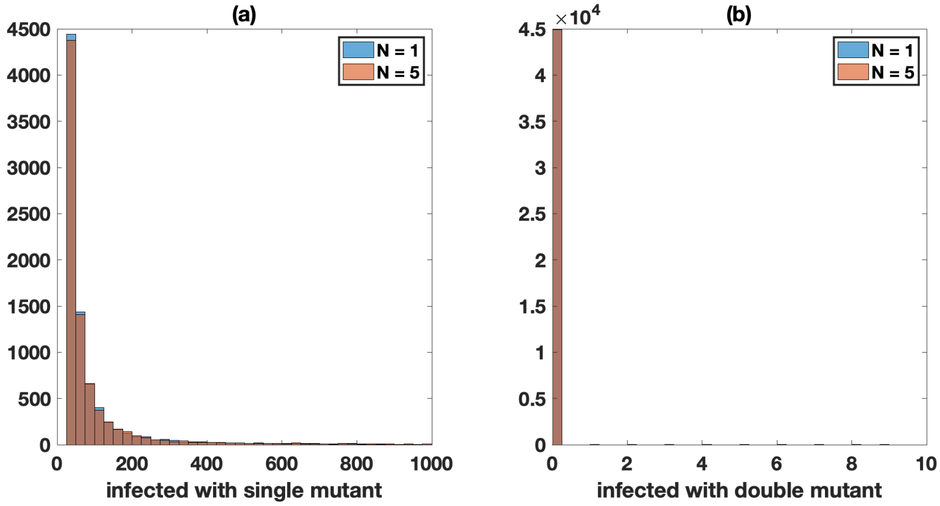

Figure S7: Neutral mutant evolution in the absence of synaptic transmission, comparing simulations with single infection only ($N = 1$, blue) and in the presence of multiple infection ($N = 5$, red). For both panels, the Kolmogorov-Smirnov test between the two distributions suggests that they are not statistically different. (a) Number of cells infected with one of the single mutant strains. (b) Number of cells infected with the double mutant strain. Distributions represent $4.5 \times 10^4$ hybrid simulations with size threshold $\mathcal{M} = 50$. Simulations are stopped at a low viral load ($10^5$ cells). The other parameters are as in main text Figure 2 ($F_{\text{wild-type}} = 1,\ F_{\text{mutant}} = 1,\ \mu = 3 \times 10^{-5},\ \lambda = 1.59 \times 10^7,\ \beta = 3.60 \times 10^{-9},\ \gamma = 0,\ a = 0.45$, and $d = 0.016$).

virus loads. This figure should be compared with Figure 2 of the main text where the mutant numbers were measured at a higher population size, and a significant increase in the number of mutants was observed in the presence of multiple infection.

Figure S8 is similar to Figure 2 of the main text, but it studies the numbers of cells infected with disadvantageous (advantageous) mutants with and without multiple infection, in the presence of only free virus transmission. As in the main text Figure 2, the number of cells was recorded near peak infection, and the maximum multiplicity parameter was $N = 1$ for simulations with single infection only, and $N = 11$ for the model including multiple infection. For non-neutral mutants we observe the same trends discussed in the main text, that is multiple infection results in an increased number of cells infected with the mutant as well as increased variation in the number of mutants. In the case of a disadvantageous single mutant with a 10% disadvantage, we observe a 2.1-fold increase in the mean number of cells infected with the mutant, and a 3.3-fold increase in the standard deviation when multiple infection is included. In the case of an advantageous single mutant with a 10% advantage, we observe a 1.9-fold increase in the mean number of cells infected with the mutant, and a 1.5-fold increase in the standard deviation when multiple infection is included.

Multiple infection also increases the prevalence of neutral triple mutants. Figure S9 compares the number of triple mutants with and without multiple infection near peak infection, again for only free virus transmission. The number of triple mutants is small, but the probability to have at least one cell infected by a triple-mutant is much higher when multiple infection is allowed (3.3% under multiple infection, which is about 1.7 times higher than the 1.9% under single infection).

With multiple infection, we can also include synaptic transmission. During synaptic cell-to-cell transmission, if the fitness of all strains is 1, we assume that $S \geq 1$ viruses are always transferred to the target cell per infection event. During an infection event, upon selection and infection of each virus, the chosen viral copy has a probability to mutate. This greatly increases the ability of synaptic transmission to generate mutant strains of virus. To see this, consider infection from a

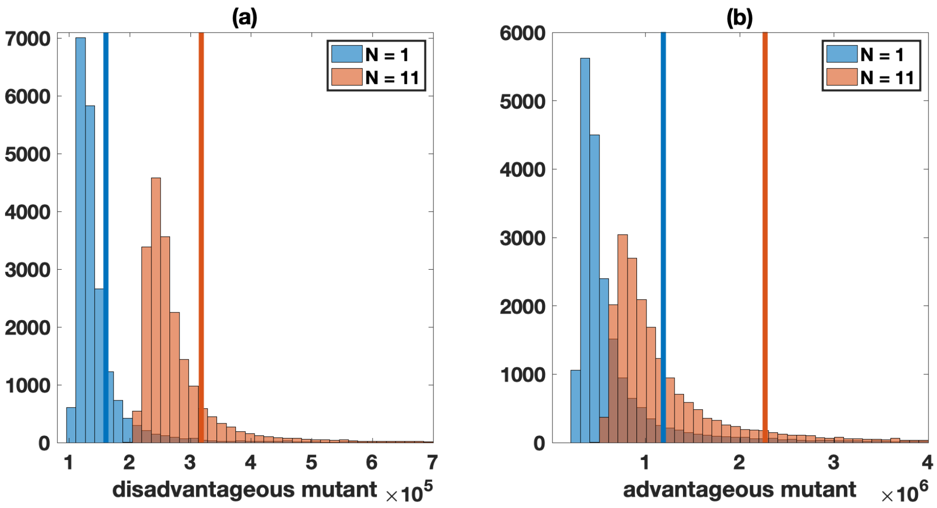

Figure S8: Non-neutral (disadvantageous and advantageous) mutant evolution in the absence of synaptic transmission, comparing simulations with single infection only ($N = 1$, blue) and in the presence of multiple infection ($N = 11$, red). The mean values are shown by vertical lines. For both panels, the Kolmogorov-Smirnov test between the two distributions gives a $p$-value less than $10^{-6}$. (a) Disadvantageous mutant; here $F_{\text{mutant}} = 0.81$. The average under single infection only is approximately $1.6 \times 10^5$ and for multiple infection is approximately $3.2 \times 10^5$. (b) Advantageous mutant with interference; here $F_{\text{mutant}} = 0.99$. The average for single infection is approximately $1.2 \times 10^6$ and for multiple infection is approximately $2.3 \times 10^6$. Histograms represent $2 \times 10^4$ hybrid simulations with size threshold $\mathcal{M} = 50$. Simulations are stopped when the infected cell population is close to peak infection ($6 \times 10^8$ cells). The other parameters are $F_{\text{wild-type}} = 0.9$, $\mu = 3 \times 10^{-5}$, $\lambda = 1.59 \times 10^7$, $\beta = 4 \times 10^{-9}$, $\gamma = 0$, $a = 0.45$, and $d = 0.016$.

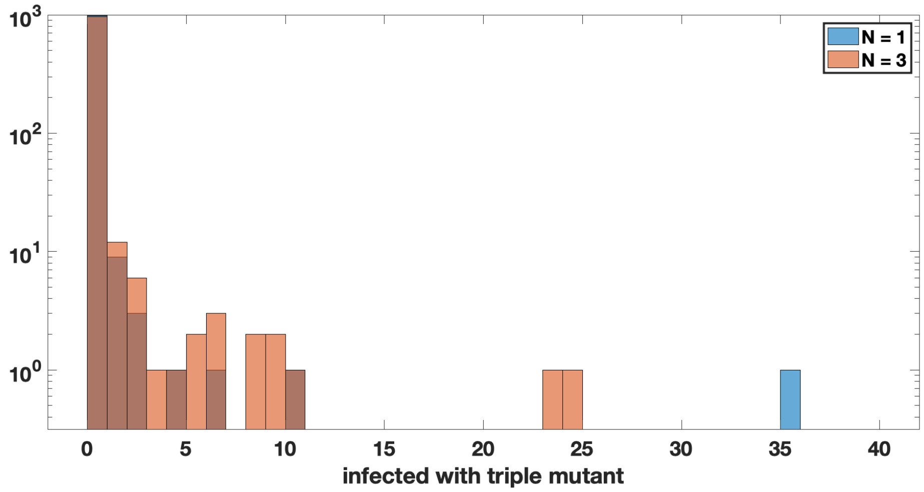

Figure S9: Presence of the triple mutant strain in the absence of synaptic transmission, comparing simulations with single infection only ($N = 1$, blue) and in the presence of multiple infection ($N = 3$, red). The probability to have at least one cell infected by a triple-mutant is 3.3% under multiple infection, which is about 1.7 times higher than that under single infection (1.9%). This result is significant with $p = 2.5 \times 10^{-3}$ by the Z-test, with $2.7 \times 10^4$ runs under single infection and $1.9 \times 10^3$ runs under multiple infection. Histograms represent $10^3$ hybrid simulations with size threshold $\mathcal{M} = 50$. Simulations are stopped when the infected cell population is close to peak infection ($6 \times 10^8$ cells). All strains are neutral ($F = 1$) and all other parameters are $\mu = 3 \times 10^{-5}$, $\lambda = 1.59 \times 10^7$, $\beta = 3.60 \times 10^{-9}$, $\gamma = 0$, $a = 0.45$, and $d = 0.016$.

cell that is infected only by the wild-type. During a free virus transmission event, there is only one chance for mutation into a mutant strain. However, during a synaptic transmission event, there are $S$ chances for mutation into a mutant strain.

If the fitness $F$ of any strain is less than 1, then there is also a chance that an unsuccessful infection event occurs. In particular, the probability of a completely unsuccessful infection event in which no viral copies are transferred during a free virus transmission event is $1-F$, and in the case of synaptic transmission it is $(1-F)^S$. Therefore, the rate at which an infection grows is influenced by the relative contribution of free virus versus synaptic transmission (and increases monotonically with increasing contribution of synaptic transmission).

Including synaptic transmission results in increased multiplicity of infection. Therefore, we need to adjust the maximum infection multiplicity used in the model, to avoid spurious accumulation of cell numbers at the end of the infection cascade. For our model, for $R_0 = 8$, $S = 3$, and 100% synaptic transmission, most cells do not become infected with more than 26 copies of virus by the time of peak infection. Therefore, we set $N = 25$ for all simulations that include synaptic transmission.

Figure S10 shows the number and distributions of cells infected with a neutral mutant ($F_{\text{wild-type}} = F_{\text{mutant}} = 1$, so infection always goes through) near peak infection for multiple combinations of free virus and synaptic transmission. As synaptic transmission is better at generating mutants, larger contributions of synaptic transmission result in larger mutant numbers and the number of mutants decreases monotonically with increasing contribution of free virus transmission.

Figure S11 should be compared with Figure 3 of the main text. For these simulations, in order to explicitly analyze the effect of multiple infection and coinfection, we initially begin with a single infected cell, which is coinfected with a single copy of both the wild-type and mutant virus. Further, we turn off mutation (set $\mu = 0$), so that synaptic transmission no longer results in a higher rate of mutant generation. Figure 3 of the main text studies disadvantageous (zero fitness) mutant spread in the absence of mutations, with and without complementation. Figure S11 considers advantageous mutants with and without interference. It shows the distribution of cells infected with a 10% advantageous mutant with and without interference for either 100% free virus or 100% synaptic transmission.

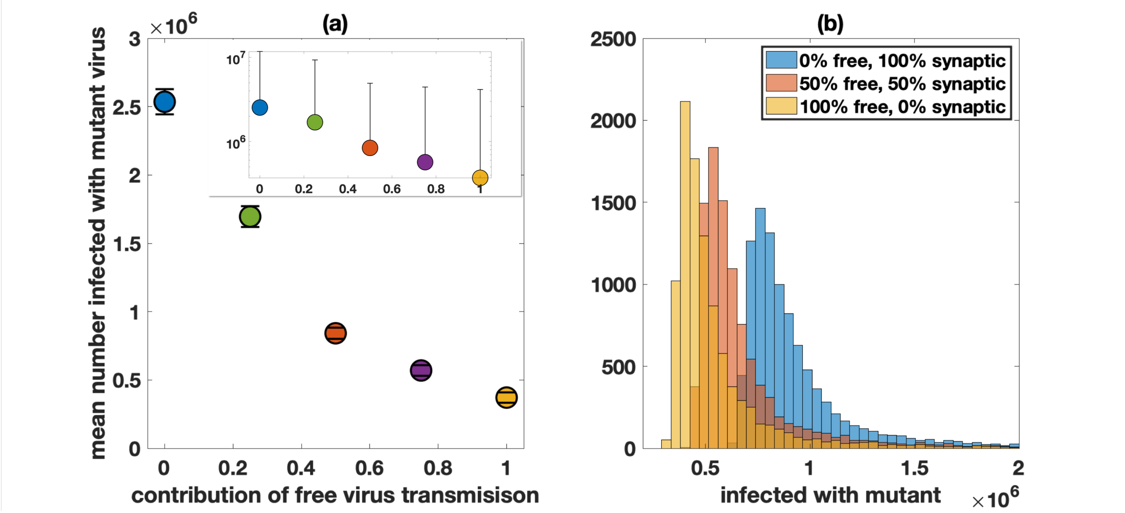

Figure S10: Neutral mutant evolution in the presence of synaptic transmission, comparing simulations with different combinations of free virus and synaptic transmission (single mutation only). (a) Number of cells infected with the mutant strain under different transmission strategies. The horizontal axis is the percent contribution of free virus transmission. Standard error bars are shown in the main figure, and standard deviation bars are shown in the inset. (b) Histograms for the number of cells infected with a single mutant for the different strategies, representing $10^4$ hybrid simulations with size threshold $\mathcal{M} = 50$. Simulations are stopped when the infected cell population is close to peak infection ($6 \times 10^8$ cells) and simulations where no infection is established are thrown out. Here $F_{\text{wild-type}} = 1$, $F_{\text{mutant}} = 1$, $S = 3$, $N = 25$, $\beta + \gamma = c = 3.6 \times 10^{-9}$, and the other parameters are $\mu = 3 \times 10^{-5}$, $\lambda = 1.59 \times 10^7$, $a = 0.45$, and $d = 0.016$.

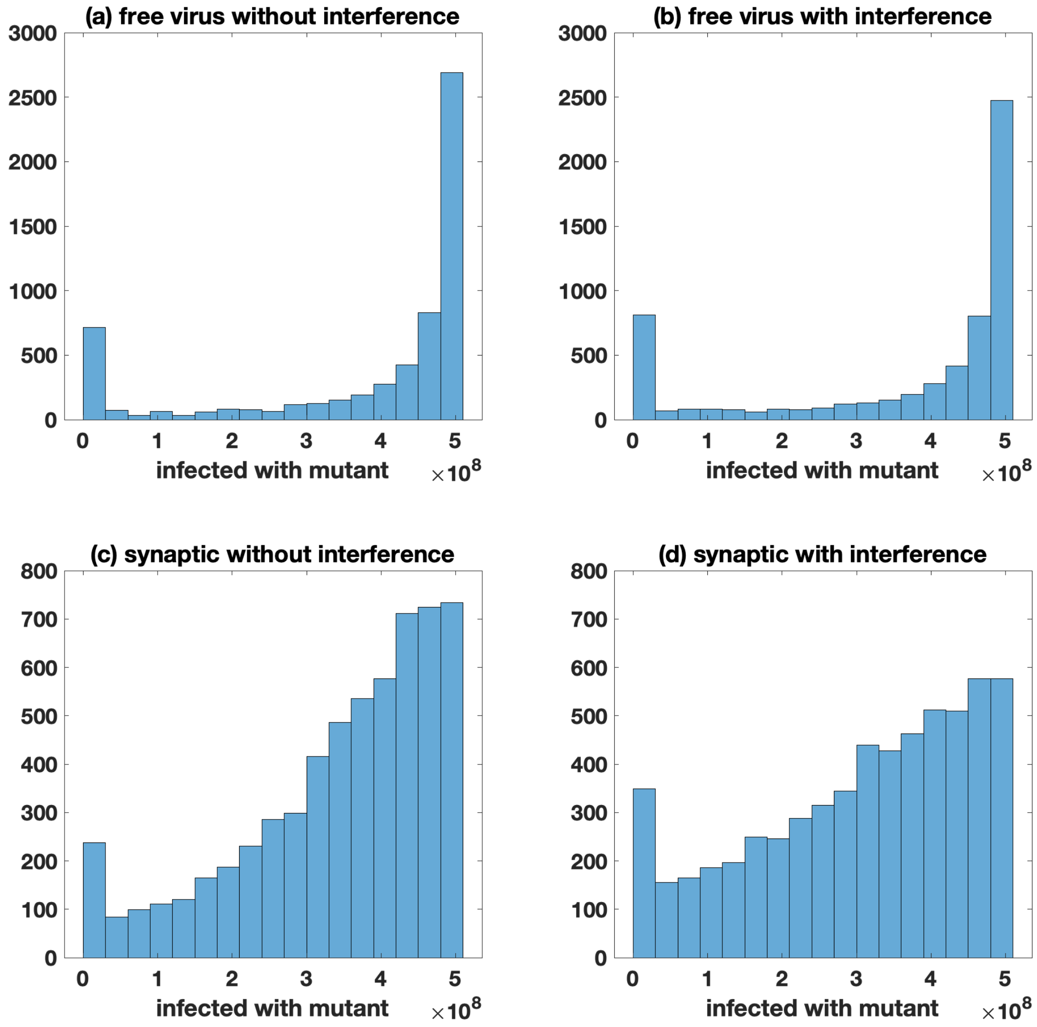

Figure S11: 10% advantageous mutant ($F_{\text{wild-type}} = 0.9, F_{\text{mutant}} = 0.99$), comparing the effect of interference and free virus versus synaptic transmission near peak infection, in the absence of mutations ($\mu = 0$). All simulations start with a single infected cell coinfected with a single copy of both the wild-type and mutant. Panels (a)-(b) represent only free virus transmission ($\beta = 3.60 \times 10^{-9},\ \gamma = 0,\ N = 11$), whereas panels (c)-(d) represent only synaptic transmission ($\beta = 0,\ \gamma = 3.60 \times 10^{-9},\ N = 25$). The Kolmogorov-Smirnov test between panels (a)-(b) and panels (c)-(d) gives a $p$-value less than $10^{-6}$. (a) Only free virus transmission without interference. The average number of cells infected with the mutant is $3.8 \times 10^8$. (b) Only free virus transmission with interference. The average number of cells infected with the mutant is $3.6 \times 10^8$. (c) Only synaptic transmission without interference. The average number of cells infected with the mutant is $3.4 \times 10^8$. (d) Only synaptic transmission with interference. The average number of cells infected with the mutant is $3.1 \times 10^8$. Histograms represent $6 \times 10^3$ hybrid simulations with size threshold $\mathcal{M} = 50$. Simulations in which infections are not established (or in the rare case a simulation does not reach the infected size threshold) are discarded; simulations are stopped when the infected cell population is close to peak infection ($5 \times 10^8$ cells). The other parameters are $\lambda = 1.59 \times 10^7,\ a = 0.45$, and $d = 0.016$.

#### References

[1] L.J.S. Allen and P. van den Driessche. Relations between deterministic and stochastic thresholds for disease extinction in continuous- and discrete-time infectious disease models. *Mathematical Biosciences*, 243(1):99 – 108, 2013.

[2] Tae-Wook Chun, Lucy Carruth, Diana Finzi, Xuefei Shen, Joseph A DiGiuseppe, Harry Taylor, Monika Hermankova, Karen Chadwick, Joseph Margolick, Thomas C Quinn, et al. Quantification of latent tissue reservoirs and total body viral load in hiv-1 infection. *Nature*, 387(6629):183–188, 1997.

[3] Tae-Wook Chun, Lucy Carruth, Diana Finzi, Xuefei Shen, Joseph A. DiGiuseppe, Harry Taylor, Monika Hermankova, Karen Chadwick, Joseph Margolick, Thomas C. Quinn, Yen-Hong Kuo, Ronald Brookmeyer, Martha A. Zeiger, Patricia Barditch-Crovo, and Robert F. Siliciano. Hiv recombination: what is the impact on antiretroviral therapy? *Nature*, 387(6629):183–188, 1997.

[4] Lina Josefsson, Martin S King, Barbro Makitalo, Johan Brännström, Wei Shao, Frank Maldarelli, Mary F Kearney, Wei-Shau Hu, Jianbo Chen, Hans Gaines, et al. Majority of CD4+ t cells from peripheral blood of HIV-1–infected individuals contain only one HIV DNA molecule. *Proceedings of the National Academy of Sciences*, 108(27):11199–11204, 2011.

[5] Lina Josefsson, Sarah Palmer, Nuno R Faria, Philippe Lemey, Joseph Casazza, David Ambrozak, Mary Kearney, Wei Shao, Shyamasundaran Kottilil, Michael Sneller, et al. Single cell analysis of lymph node tissue from HIV-1 infected patients reveals that the majority of CD4+ T-cells contain one HIV-1 DNA molecule. *PLoS pathogens*, 9(6):e1003432, 2013.

[6] Andreas Jung, Reinhard Maier, Jean-Pierre Vartanian, Gennady Bocharov, Volker Jung, Ulrike Fischer, Eckart Meese, Simon Wain-Hobson, and Andreas Meyerhans. Recombination: Multiply infected spleen cells in hiv patients. *Nature*, 418(6894):144, 2002.

[7] Natalia L Komarova and Dominik Wodarz. Virus dynamics in the presence of synaptic transmission. *Mathematical biosciences*, 242(2):161–171, 2013.

[8] Martin A Nowak, Alun L Lloyd, Gabriela M Vasquez, Theresa A Wiltrout, Linda M Wahl, Norbert Bischofberger, Jon Williams, Audrey Kinter, Anthony S Fauci, Vanessa M Hirsch, et al. Viral dynamics of primary viremia and antiretroviral therapy in simian immunodeficiency virus infection. *Journal of virology*, 71(10):7518–7525, 1997.

[9] John E. Pearson, Paul Krapivsky, and Alan S. Perelson. Stochastic theory of early viral infection: Continuous versus burst production of virions. *PLOS Computational Biology*, 7(2):1–17, 02 2011.

[10] Alan S Perelson, Avidan U Neumann, Martin Markowitz, John M Leonard, and David D Ho. Hiv-1 dynamics in vivo: virion clearance rate, infected cell life-span, and viral generation time. *Science*, 271(5255):1582–1586, 1996.

[11] Ruy M Ribeiro, Li Qin, Leslie L Chavez, Dongfeng Li, Steven G Self, and Alan S Perelson. Estimation of the initial viral growth rate and basic reproductive number during acute HIV-1 infection. *Journal of virology*, 84(12):6096–6102, 2010.

[12] Ignacio A. Rodriguez-Brenes, Natalia L. Komarova, and Dominik Wodarz. The role of telomere shortening in carcinogenesis: A hybrid stochastic-deterministic approach. *Journal of Theoretical Biology*, 460:144 – 152, 2019.

[13] Alex Sigal, Jocelyn T Kim, Alejandro B Balazs, Erez Dekel, Avi Mayo, Ron Milo, and David Baltimore. Cell-to-cell spread of HIV permits ongoing replication despite antiretroviral therapy. *Nature*, 477(7362):95, 2011.

[14] P. Whittle. The outcome of a stochastic epidemic—a note on Bailey's paper. *Biometrika*, 42(1-2):116–122, 06 1955.

[15] Dominik Wodarz, David N Levy, and Natalia L Komarova. Multiple infection of cells changes the dynamics of basic viral evolutionary processes. *Evolution letters*, 3(1):104–115, 12 2018.
